# Genetic differentiation and intrinsic genomic features explain variation in recombination hotspots among cocoa tree populations

**DOI:** 10.1101/482299

**Authors:** Enrique J. Schwarzkopf, Juan C. Motamayor, Omar E. Cornejo

## Abstract

Our study investigates the possible drivers of recombination hotspots in *Theobroma cacao* using ten genetically differentiated populations. By comparing recombination patterns between multiple populations, we obtain a novel view of recombination at the population-divergence timescale. For each population, a fine-scale recombination map was generated using the coalescent with a standard method based on linkage disequilibrium (LD). These maps revealed higher recombination rates in a domesticated population and a population that has undergone a recent bottleneck. We inferred hotspots of recombination for each population and find that the genomic locations of hotspots correlate with genetic differentiation between populations (F_ST_). We used randomization approaches to generate appropriate null models to understand the association between hotspots of recombination and both DNA sequence motifs and genomic features. We found that hotspot regions contained fewer known retroelement sequences than expected and were overrepresented near transcription start and termination sites. Our findings indicate that recombination hotspots are evolving in a way that is consistent with genetic differentiation but are also preferentially driven to near coding regions. We illustrate that, consistent with predictions in plant domestication, the recombination rate of the domesticated population is orders of magnitude higher than that of other populations. More importantly, we find two fixed mutations in the domesticated population’s FIGL1 protein. FIGL1 has been shown to increase recombination rates in *Arabidopsis* by several orders of magnitude, suggesting a possible mechanism for the observed increased recombination rate in the domesticated population.

## Introduction

Genetic recombination is an important source of genome-wide genetic variation fundamental for evolutionary forces like selection and genetic drift to act. Selection and drift contribute to a loss of variation, which means that in the absence of forces that maintain variation along the genome, populations would be incapable of evolving over prolonged periods of time. Recombination rearranges genetic material onto different backgrounds generating a larger set of haplotype combinations on which selection can act, reducing the magnitude of Hill-Robertson interference (Felsenstein 1974). Different regimes of recombination can strongly influence how efficient selection is at purging deleterious mutations and increasing the frequency of beneficial mutations in the population (Felsenstein 1974).

Studies in a wide range of species have shown that recombination rates are not uniform along the genome and general patterns of variation have been described (Begun and Aquadro 1992, Akhunov et al. 2003, Wu et al. 2003, Anderson et al. 2004, McVean et al. 2004, Mézard 2006, Kim et al. 2007, Gore et al. 2009, Schnable et al. 2009, Branca et al. 2011,Paape et al. 2012). One pattern that has been observed in multiple species is the reduced recombination rate in centromeric regions of the chromosomes and the progressive increase of recombination rates as the physical distance from the telomeres decreases (Begun and Aquadro 1992, Akhunov et al. 2003, Wu et al. 2003, Anderson et al. 2004, Gore et al. 2009, Schnable et al. 2009). This pattern has also been shown to arise in simulation studies (e.g. Mackiewicz et al. 2010). Another interesting pattern that has been observed is that of regions with unusually high rates of recombination spread throughout chromosomes: recombination hotspots (McVean et al. 2004, Brunschwig et al. 2012, Paape et al. 2012, Hellsten et al. 2013, Stevison et al. 2016, Shanfelter et al. 2018). The importance of recombination hotspots lies in their ability to shuffle genetic variation at higher rates than the rest of the genome, profoundly impacting the dynamics of selection for or against specific mutations (Felsenstein 1974). In this study, we focus on locally defined recombination hotspots, requiring that their recombination rate be unusually high when compared to neighboring regions.

A variety of genomic features have been identified as being associated with regions of high recombination. Recombination hotspots have been linked to transcriptional start sites (TSSs) and transcriptional termination sites (TTSs) in *Arabidopsis thaliana*, *Taeniopygia guttata*, *Poephila acuticauda*, and humans (Myers et al. 2005, Choi et al. 2013, Singhal et al. 2015). In *Mimulus guttatus* hotspots were found to be associated with cpg islands (short segments of cytosine and guanine rich DNA, associated with promoter regions) (Hellsten et al. 2013). CpG islands were also associated with increased recombination rates in humans and chimpanzees (Auton et al. 2012). These patterns point to recombination occurring frequently near, but not within, coding regions. The formation of chiasmata is important for the proper disjunction of chromosomes during meiosis (Martinez-Perez et al. 2008), but repeated double-strand breaks can lead to an increased mutation rate (Rodgers and McVey 2015). In coding regions in particular, this excess mutation rate can have a high evolutionary cost, due to the likelihood of novel deleterious mutations being higher than that of beneficial ones (Haldane 1937, Crow and Kimura 1970, Wloch et al. 2001, Sanjuán et al. 2004, Eyre-Walker and Keightley 2007). Recombination hotspots have also been found to be correlated with particular DNA sequence motifs. In some mammals, including *Mus musculus* (Brunschwig et al. 2012) and apes (Auton et al. 2012, Stevison et al. 2016) binding sites for PRDM9, a histone trimethylase with a DNA zinc-finger binding domain, have been found to correlate with recombination hotspots. In *A. Thaliana*, proteins that limit overall recombination rate have been identified, leading to a genome-wide increase in recombination rate in knockout mutants (Fernandes et al. 2018b). However, these *Arabidopsis* proteins have not been shown to direct recombination to particular regions and are therefore not expected to affect the location of recombination hotspots.

The dynamics of recombination hotspots shared between related species or populations of the same species have been investigated in apes, yielding varying results. Hinch et al. (2011) found that, at finer scales, the genetic maps of European and African human populations were significantly different. They also found that, when looking at hotspots in the major histocompatibility complex, the African populations showed a hotspot that was not present in Europeans, but all European hotspots were found in African populations (Hinch et al. 2011). Recent work on recombination in apes found little correlation of recombination rates in orthologous hotspot regions when looking between species, but a strong correlation when comparing between two populations of the same species (Stevison et al. 2016). Other studies have also found very little sharing of hotspots between humans and chimpanzees (Ptak et al. 2005, Winckler et al. 2005). Additionally, the dynamic of changing hotspot locations observed in humans and other apes has been observed in simulations (Mackiewicz et al. 2013). The disparity of empirical results regarding hotspots shared between related populations suggest that further work is required to disentangle the relationship between demographics and shared hotspots.

The identification of ten genetically differentiated populations of the cocoa tree, *Theobroma cacao*, (Motamayor et al. 2008, Cornejo et al. 2017) can be leveraged to study population-level dynamics of recombination patterns. The ten *T. Cacao* populations originate from different regions of South and Central America, and include one fully domesticated population (Criollo), used in the production of fine chocolate, and nine wilder, more resilient populations which generate higher cocoa yield than the Criollo variety (Fig. S1) (Motamayor et al. 2008, Henderson et al. 2007, Cornejo et al 2017). These ten populations have been shown to have strong signatures of differentiation between them (F_ST_ values ranging from 0.16 to 0.65) and they separate into clear clusters of ancestry (Cornejo et al. 2017). During domestication, recombination plays an important role in the segregation of traits, and for this reason it has been hypothesized that recombination rates will increase during the process of domestication (Moyers et al. 2018). Domestication can be a rapid process and there is theoretical evidence for the increase of recombination rates during periods of rapid evolutionary change (Otto and Barton 1997). Empirical evidence for this prediction has been shown in a limited number of herbaceous plant species with short generation times (Ross-Ibarra 2004). It is not clear if plant species with longer generation times are also expected to experience increased recombination rates, and it is also unclear what mechanisms could explain these differences. One possible explanation for differences in recombination rates between wild and domesticated populations is polymorphism in genes like those previously demonstrated to suppress recombination in *Arabidopsis thaliana* (Girard et al. 2015, Fernandes et al. 2018a). The differences in recombination rates between wild and domesticated populations is just one of the possible questions that can be touched on with this system.

The ten populations of *T. cacao* also allow us to compare the locations of hotspots between them, potentially contributing to the understanding of hotspot turnover at the population-divergence timescale. These comparisons can also contribute to our understanding of how demographics impact the turnover of recombination hotspot locations. *T. cacao* is unique in this case for being a long-lived organism with no known driver of recombination hotspots (e.g. PRDM9). What contributes to the location of recombination hotspots in such a species is, of course, contingent on our being able to detect recombination hotspots in the different populations of *T. cacao*.

In order to locate recombination hotspots for *T. Cacao* populations, we must first obtain fine-scale recombination maps for each population, which we did using an LD-based method. Fine-scale, LD-based recombination maps have been constructed for a number of plant models (Paape et al. 2012, Choi et al. 2013, Hellsten et al. 2013), identifying a variety of features correlated to recombination rate. Unlike these model plants with short generation times, *T. Cacao* is a perennial woody plant with a five-year generation time (Henderson et al. 2007). The size and long generation time of *T. Cacao* makes direct measurements of recombination impractical. However, historical recombination can be estimated for *T. Cacao* using coalescent based methods (Auton and McVean 2007). Theoretical studies have shown that population structure can generate artificially inflated measures of LD (Li and Nei 1974, Ohta 1982), which would be detrimental to our estimates of recombination. For this reason, recombination maps were constructed independently for each population. For each population we aim to describe the relationship between recombination hotspots and a variety of evolutionary and genomic factors.

We used an LD-based method to estimate recombination rates for ten populations of *T. Cacao*, which we then analyzed with a maximum likelihood statistical framework to infer the location of recombination hotspots. The locations of hotspots were compared across populations and a novel resampling scheme tailored to the genomic architecture of *T. Cacao* was used to generate null assumptions for the distribution of hotspots along the genome. These null distributions were used to identify differential representation of known DNA sequence motifs in ubiquitous recombination hotspots, and of overlap between recombination hotspots and genomic traits for each population. The re-sampling schemes used to identify these associations are novel in the context of this work and were designed to take into account the size and distribution of elements in the genome. In this work we aimed to answer the following questions: (i) How are recombination rates distributed within 10 highly differentiated populations of *T. Cacao*, and how do they compare to each other? (ii) How are hotspots distributed along the genome of each of the ten populations of *T. Cacao*, and can these distributions be explained by patterns of population genetic differentiation? (iii) Are there identifiable DNA sequence motifs that are associated with the location of recombination hotspots along the *T. Cacao* genome? (iv) Are there genomic features (e.g. TSSs, TTSs, exons, introns) consistently associated with recombination hotspot locations across *T. Cacao* populations? Our findings suggest that recombination hotspot locations generally follow patterns of diversification between populations, while also having a strong tendency to occur close to TSSs and TTSs. Moreover, we find a strong negative association between the occurrence of recombination hotspots and the presence of retroelements.

## Results

### Comparing recombination rates between populations

Populations show a mean recombination rate (*r*) between 2.1 and 525 cm/Mb (Table 1), with a variety of distributions (Fig. S2). We observe a higher mean than median *r* indicating that extreme high values are present for all populations. The extreme recombination rate values affect the mean, driving it to values consistently higher than the median. The pattern of recombination rates along the genome varied between populations, as can be seen in the comparison of the Nanay and Purus third chromosome (Fig. 1). Purus appears to have a higher average recombination rate than Nanay for chromosome three. More specifically, particular regions of the chromosome present peaks in one population that are absent in the other. A similar patter can also be observed for the density of recombination hotspots, e.g. Purus presenting a high density of hotspots in certain regions that is not observed in Nanay. The median 95% probability interval for recombination rate across the genome for each population was found to be several orders of magnitude larger than the uncertainty per site, estimated as the median 95% Credibility Interval of the trace for each position in the genome for that population (Table S1).

**Table 1.** Recombination rates in *ρ = 4Ner* (in Morgans per base) and *r* (in cM/Mb) for all ten *T. cacao* populations. The *Ne* (from Cornejo et al., 2018) used to transform *ρ* to *r* for each population is also reported, as are the lower and upper bounds of a 95% confidence interval for mean *r*.

| Population | Mean $4N_e r$ | Mean $N_e$ | Mean $r$ (cM/Mb) | Median $r$ (cM/Mb) | L bound (Mean $r$) | U bound (Mean $r$) |
| --- | --- | --- | --- | --- | --- | --- |
| Amelonado | 1.58e-03 | 15,744 | 2.51 | 2.40e-04 | 2.48 | 2.54 |
| Contamana | 8.53e-03 | 61,102 | 3.49 | 4.92e-01 | 3.48 | 3.50 |
| Criollo | 1.46e-04 | 695 | 525 | 427 | 523 | 527 |
| Curaray | 1.04e-04 | 58,213 | 4.45 | 1.78 | 4.44 | 4.46 |
| Guianna | 8.66e-03 | 4,651 | 46.5 | 7.74e-01 | 46.3 | 46.7 |
| Iquitos | 4.23e-03 | 49,984 | 2.11 | 5.88e-04 | 2.10 | 2.12 |
| Maranon | 4.09e-03 | 34,037 | 3.01 | 1.64e-03 | 2.99 | 3.02 |
| Nacional | 4.66e-03 | 26,060 | 4.47 | 9.76e-03 | 4.44 | 4.49 |
| Nanay | 6.82e-03 | 42,429 | 4.02 | 1.51e-02 | 4.00 | 4.04 |
| Purus | 5.95e-03 | 17,357 | 8.57 | 7.74e-01 | 8.54 | 8.60 |

**Figure 1.**
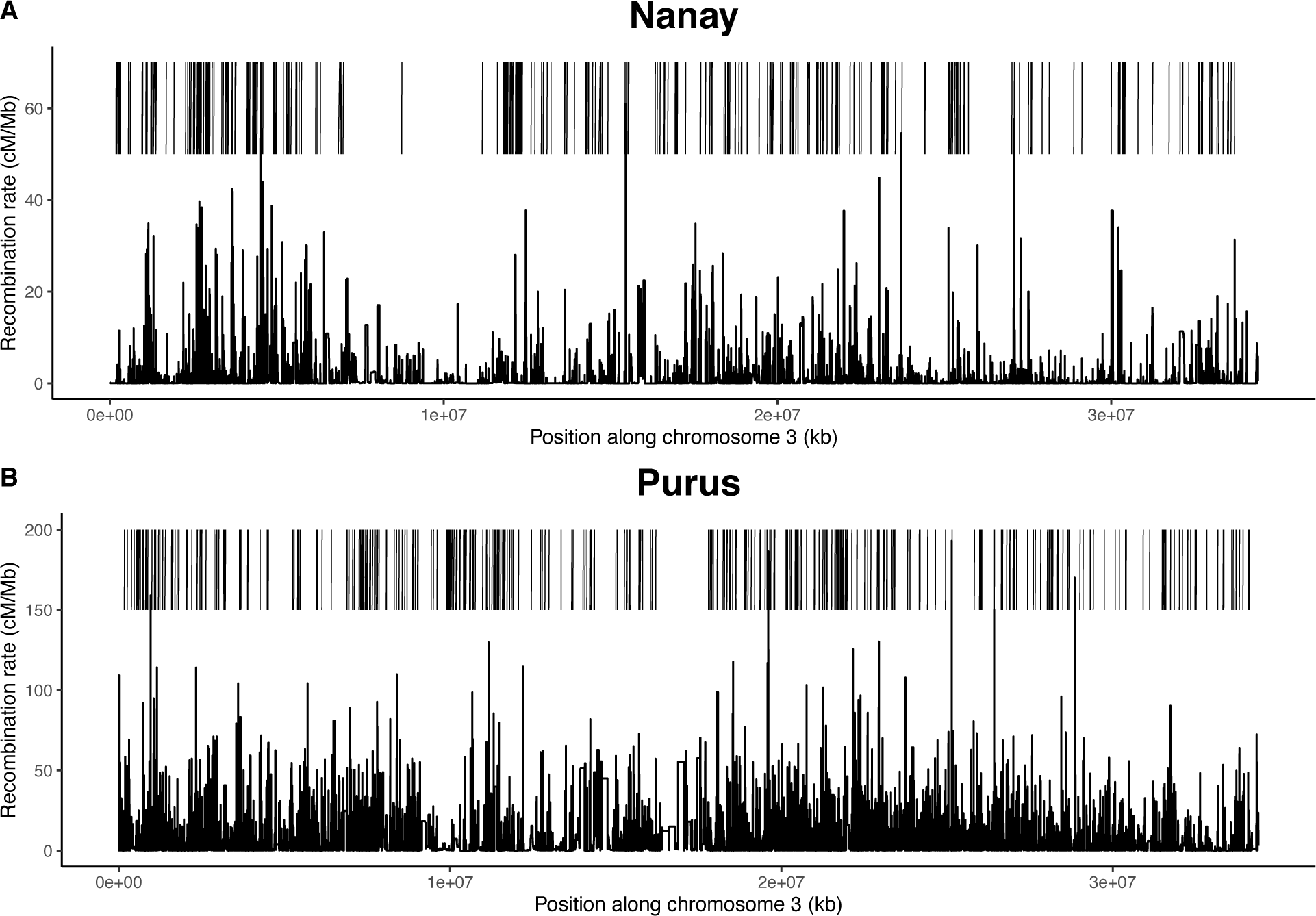
The third chromosomes of the Nanay (A) and Purus (B) populations were selected to exemplify the differences between populations in recombination rates (*r*) and recombination hotspot locations (bars above rates).

Overall, the mean recombination rate for most of the populations is similar to that found for *Arabidopsis thaliana* using LDhat, when using θ=0.1 (Choi et al. 2013) (Table 1). The LDhat estimates using θ=0.001 were slightly higher than the estimates using θ=0.1 for each population. We chose to proceed with analyses using the results from the θ=0.1 since more of the mean population recombination rates fell within the range of values identified in plants (Stapley et al. 2016) (Table S2).

In order to compare the average recombination rates of the different populations, a Wilcoxon signed-rank test was performed for every pair of populations. The only pair that did not show a significant difference in mean recombination rate was that of Nacional and Nanay (*p=0.3*). All other pairwise comparisons were highly significant (*p<2e-16*).

Two populations, Guianna and Criollo, have a higher average recombination rate than the other populations by one and two orders of magnitude respectively (Table 1). Guianna and Criollo also have been estimated to have a lower effective population size (*N*_*e*_) (Cornejo et al. 2018) by one and two orders of magnitude respectively. However, there was no significant association between mean *N*_*e*_ and *r* (*p=0.1119*), indicating that, for a high enough *N*_*e*_, the ability to detect recombination events is not dictated by the effective population size. When Criollo and Guianna were excluded, the relationship was also not present (*p=0.3886*). When all populations were included, the inbreeding coefficient (*F*, from Cornejo et al. 2018) showed no significant linear association with mean *r* (*p=0.3361*). We also found no linear trend between sample size and mean *r* (*p=0.2333*).

The FIGL1 and FLIP proteins characterized by Fernandes et al. (2018a) were found to be responsible for recombination suppression in *Arabidopsis*. Plants with a FIGL1 knockout were found to increase recombination rates significantly and FLIP knockouts show increases of recombination at a much lesser extent (Fernandes et al. 2018). Therefore, we explored the possibility that missense FIGL1 and FLIP orthologs in *T. cacao* explain the between-population differences in recombination rate. We used a reciprocal BLAST search to identify the orthologs for both genes and used annotation data from Cornejo et al. (2018) to identify 15 missense mutations in FIGL1 and 18 missense mutations in FLIP (Fig. 2, Fig. S3, Table S3). We then used a generalized linear model framework to infer the impact of the 8 uncorrelated missense mutations found in the *T. cacao* FIGL1 ortholog under the assumption of a full recessive model. We find that mutations 215KK (Coeff.=426.54, p=2.41e-13), 155II (Coeff.=8.97, p=0.000231), and 291TT (Coeff.=0.47, p=0.047) significantly explain changes in the recombination rate, but all other mutations made no significant impact. The same model was run for FLIP but returned no significant coefficients (after eliminating perfectly correlated mutations with those found in FIGL1).

**Figure 2.**
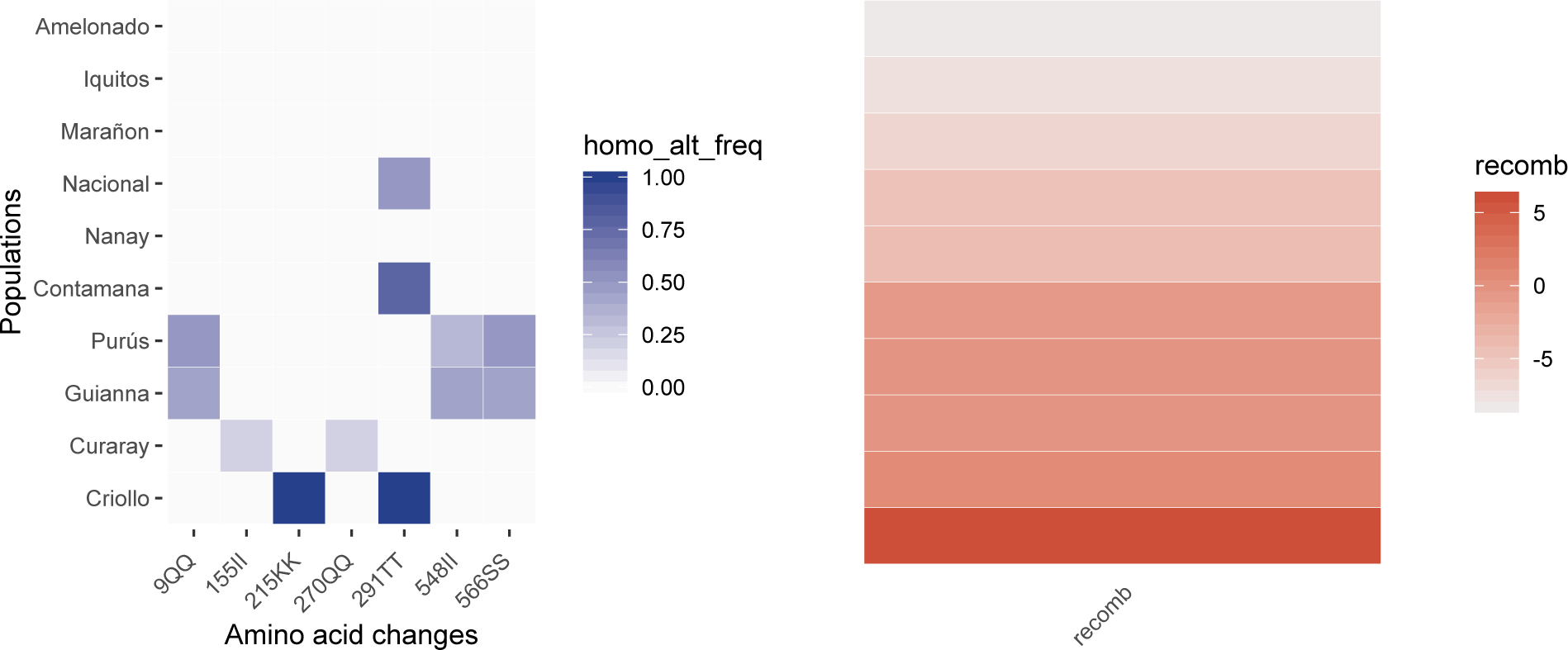
The left panel shows the frequency of individuals that are homozygous for the alternative allele of amino acid mutations in a *T. cacao* FIGL1 ortholog. Alternative allele is defined in terms of the Amelonado reference genome. The right panel shows the *log*_*e*_ transformed recombination rates (*r*). The populations are in the same order in both panels.

### Comparing recombination hotspot locations between populations

The majority (55.5%) of hotspots identified were not shared between populations. The 25 most numerous sets of hotspots are represented in Fig. 3. The nine largest of these are sets of hotspots unique to single populations. The hotspots unique to the remaining population (Criollo) formed the eleventh largest set. Effective population size (*N*_*e*_) is not a good linear predictor of the number of detected hotspots (*p=0.1489*), nor is sample size (*p=0.351*).

**Figure 3.**
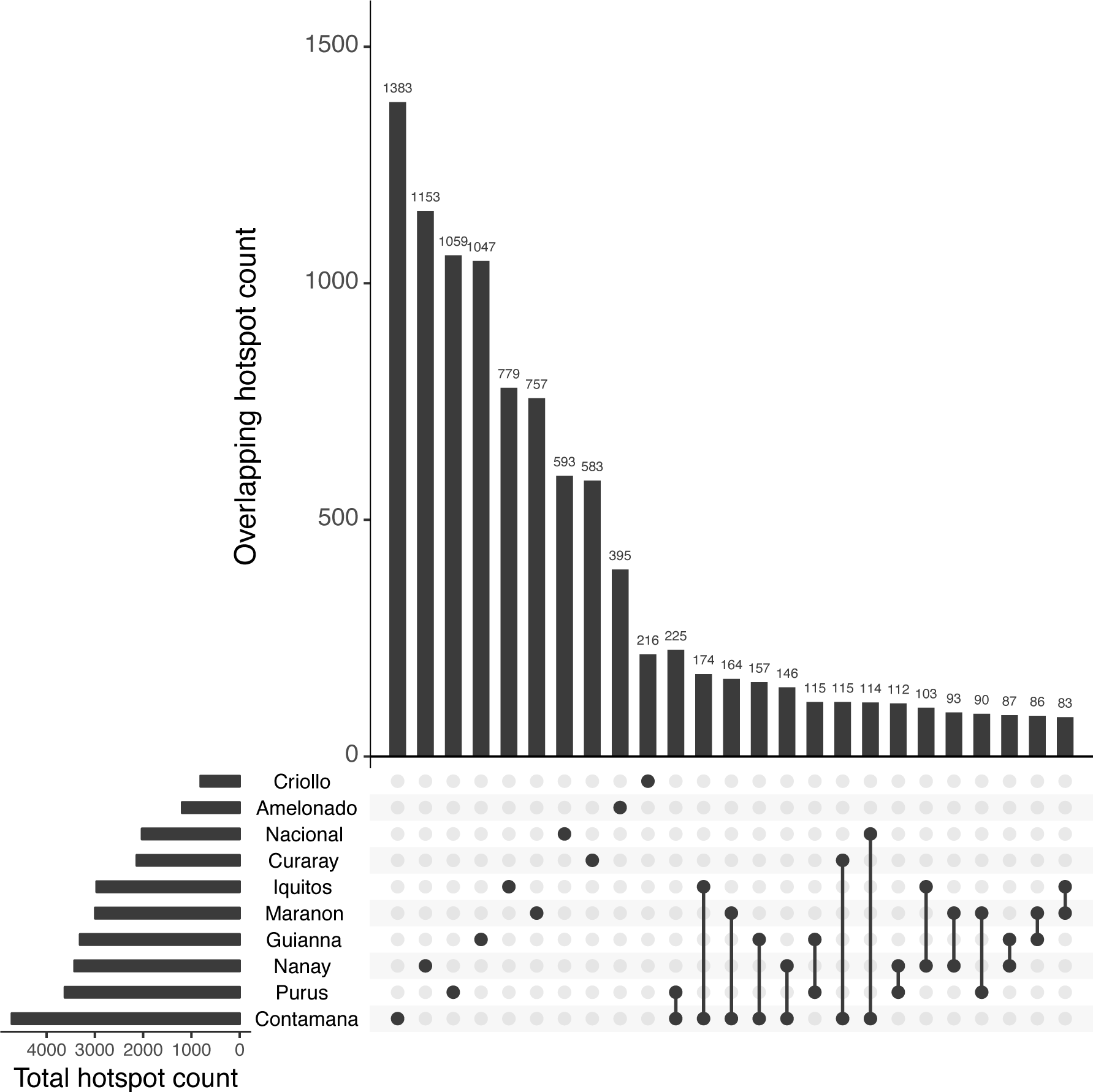
Upset plot showing number of hotspots in different subsets. Horizontal bars represent total hotspots detected in a population, each dot on the matrix indicate that the vertical bar above it is the count of hotspots unique to that population, connected dots indicate that the vertical bar above them represents hotspots shared between the populations represented by the connected dots. The 25 largest subsets are shown.

The recombination rate in hotspot regions for nine of the populations was on average between 22 and 237% higher than the average recombination rate of the genome. The exception was Guianna, which only showed an approximately 1% increase in average recombination rate in hotspots regions when compared to that of the non-hotspot regions. For Guianna, we also compared the recombination rate inside hotspots to their surrounding regions (+/− 5kb). We found that hotspot regions had a rate ~42% higher rate than their neighboring regions. This result leads us to believe that the 1% higher average recombination rate in the Guianna hotspots when compared to the entire genome may be due to an increased ability to detect hotspots in regions of low recombination for this population. Additionally, Guianna presents unusually large hotspots (average 8.9 kb, Table S4), which points to an especially low resolution in hotspot detection for this population.

Despite the majority of hotspots not being shared between populations, we conducted pairwise Fisher’s exact tests to verify whether there was significantly more hotspot overlap than expected (if hotspots were randomly distributed along the genome) between populations. For most pairs of populations, we found significantly more hotspot overlap than expected (Table 2). There were three comparisons that did not show significantly more overlap than expected: Amelonado-Nacional, Amelonado-Purus, and Criollo-Nacional. A Mantel test comparing distances between populations based on shared hotspots and F_ST_ values between populations resulted in a significant correlation between them (*r=0.66*, *p=0.002*).

**Table 2.** Fisher’s exact test p-values for pairwise comparisons of recombination hotspot locations between populations of *T. cacao*. We conducted 45 comparisons, corresponding to a Bonferroni correction cutoff value of *α=0.0011*.

| Population | Ame | Con | Cri | Cur | Gui | Iqu | Mar | Nac | Nan |
| --- | --- | --- | --- | --- | --- | --- | --- | --- | --- |
| Amelonado | - | - | - | - | - | - | - | - | - |
| Contamana | <2e-07 | - | - | - | - | - | - | - | - |
| Criollo | <9e-05 | <5e-13 | - | - | - | - | - | - | - |
| Curaray | <3e-05 | <3e-37 | <5e-08 | - | - | - | - | - | - |
| Guianna | <3e-06 | <1e-37 | <7e-07 | <4e-20 | - | - | - | - | - |
| Iquitos | <4e-08 | <6e-87 | <2e-11 | <3e-16 | <2e-29 | - | - | - | - |
| Maranon | <6e-13 | <7e-77 | <2e-11 | <2e-20 | <5e-33 | <4e-64 | - | - | - |
| Nacional | 0.0015 | <2e-43 | 0.0212 | <7e-14 | <3e-06 | <6e-14 | <3e-13 | - | - |
| Nanay | 0.0004 | <2e-44 | <9e-11 | <4e-16 | <2e-21 | <3e-39 | <2e-38 | <9e-06 | - |
| Purus | 0.1782 | <4e-117 | <2e-05 | <2e-29 | <1e-33 | <2e-39 | <8e-43 | <6e-27 | <2e-21 |

To study the effects of demographic history more closely, shared hotspots were converted to dimensions of a multiple correspondence analysis and modeled along a previously constructed drift tree (Cornejo et al. 2018). Modeling the dimension as a Brownian motion was a better fit (AIC=79.4) than modeling it as an Ornstein-Uhlenbeck (OU) process (AIC=81.4), which is consistent with the small number of hotspots shared between populations. The model assuming Brownian motion is consistent with pure drift driving differentiation of a trait along a genealogy, while an OU process is consistent with a higher trait maintenance (stabilizing selection).

### Identifying DNA sequence motifs associated with the locations of recombination hotspots

We used RepeatMasker to analyze the set of recombination hotspots that were present in at least eight *T. Cacao* populations (17 total hotspots; referred to as ubiquitous hotspots), as well as the consensus set of recombination hotspots and the reference genome. In order to determine whether a particular set of DNA sequence repeats was overrepresented in ubiquitous hotspots, the percentage of DNA sequence that was identified as potentially being from retroelements or DNA transposon was compared to an empirical distribution. The percentage of observations from the distribution which were greater than the observed are reported in Table 3. While retroelements were found to be underrepresented in the ubiquitous hotspots, DNA transposons were marginally overrepresented.

**Table 3.** Percentage of DNA sequences identified as either retroelements or DNA transposons, and total interspersed repeats. Observed values for the entire *T. cacao* genome, for all recombination hotspots (HS), and ubiquitous hotspots (hotspots in the same location in at least eight different populations). Also presented are mean percentage of these sequences for 1000 simulations of hotspots equivalent in size and count as the ubiquitous set and the percentile at which the observed value for the ubiquitous set is found in the distribution of the simulated set (Sim).

| Measures | Observed %<br>ubiquitous HS | Observed %<br>all HS | Observed %<br>whole genome | Mean %<br>Sim | % Sim ><br>ubiquitous |
| --- | --- | --- | --- | --- | --- |
| Retroelements | 2.34 | 9.45 | 11.12 | 11.11 | 99.9 |
| DNA transposons | 1.94 | 1.64 | 1.10 | 1.10 | 5.4 |
| Total | 4.28 | 11.09 | 12.21 | 12.22 | 99.7 |

### Identifying genomic features associated with the location of recombination hotspots

We found an overrepresentation of recombination hotspots at transcriptional start sites (TSSs) and transcriptional termination sites (TTSs) in all ten of the *T. Cacao* populations (Table 4). The level of overrepresentation of hotstpots in particular regions was compared to a null expectation based on simulations of hotspots of the same size as the ones detected, distributed randomly along the chromosomes. For all populations, all 1000 simulations showed a lower proportion of overlap with TSSs and TTSs than the observed. In the case of exons and introns, seven populations (Contamana, Criollo, Iquitos, Maranon, Nacional, Nanay, Purus) had an observed value that was lower than all, or almost all (Purus for exons), simulations. Three of the remaining four populations (Amelonado, Curaray, and Nanay) had no clear trend in either direction (Table 4). The final population (Guianna) showed an overrepresentation of hotspots in both exons and introns.

**Table 4.** Proportion of simulated chromosomes that presented a lower number of hotspots intersecting with TSSs, TTSs, exons, and introns than the observed chromosomes. TSSs and TTSs are considered to be the 500bp upstream and downstream of transcribed regions, respectively.

| Population | TSSs<br>(500bp) | TTSs<br>(500bp) | Exons | Introns |
| --- | --- | --- | --- | --- |
| Amelonado | 1 | 1 | 0.602 | 0.527 |
| Contamana | 1 | 1 | 0 | 0 |
| Criollo | 1 | 1 | 0 | 0 |
| Curaray | 1 | 1 | 0.346 | 0.058 |
| Guianna | 1 | 1 | 1 | 1 |
| Iquitos | 1 | 1 | 0 | 0 |
| Maranon | 1 | 1 | 0 | 0 |
| Nacional | 1 | 1 | 0 | 0 |
| Nanay | 1 | 1 | 0.027 | 0.237 |
| Purus | 1 | 1 | 0.004 | 0 |

## Discussion

The set of ten *T. cacao* populations, which includes wild, long-established and recently established populations as well as a domesticated population, has provided us a unique opportunity to study differences in recombination in populations of the same species with varying evolutionary histories (Cornejo et al. 2018, Bartley et al. 2005). We also explored differences in the location of recombination hotspots between the populations and found that the conservation of hotspots between them generally mirrors their patterns of pairwise genetic differentiation. Additionally, we found that TSSs and TTSs are strongly associated with recombination hotspots in all populations, which is consistent with previous findings in plants (Paape et al. 2012, Choi et al. 2013, Hellsten et al. 2013). This factor seems to play an important role in determining the location of novel hotspots. Finally, hotspots that are shared by at least eight populations appear to be associated with DNA transposons, pointing to a potential mechanism for the maintenance of recombination hotspots at the population-divergence timescale. Understanding how recombination rates vary between genetically differentiated populations of the same species is an important step toward disentangling the role of recombination in genetic differentiation.

### Comparing recombination rates between populations

We found that the eight long-established, wild *T. cacao* populations show an average recombination rate (*r*) within the range of recombination rates measured for other suporrosids (Table 1; Stapley et al. 2016). The average recombination rate for these *T. cacao*, populations, was 4.07 cM/Mb, which is very similar to that of *Juglans regia* (4.05 cM/Mb; Zhu et al. 2015), and comparable to other woody rosids (e.g. *Populus deltoides*, 8.13 cM/Mb (Mousavi et al. 2016); *Mangifera indica*, 7.15 cM/Mb (Luo et al. 2016); *Citrus clementina*, 3.6 cM/Mb (Ollitrault et al 2012)) (Stapley et al. 2016). This places *T. cacao* on the high end of known recombination rates for its order but comfortably in the range of other long-lived, woody plants. For all ten populations, the mean recombination rate was found to be greater than the median. This is consistent with high rate outlier values; an expected result in the presence of recombination hotspots.

The other two *T. cacao* populations (Criollo, the domesticated variety and Guianna, the recently established wild population) show unusually high average recombination rates when compared to the eight long-established wild populations. Despite a small sample for some populations, including Criollo, we found no linear trend between sample size and recombination rate (Table S5). Additionally, the rates calculated for the two wild, small-sample populations (Curaray and Nacional) were consistent with those of other wild populations. This makes us confident in our estimates for the Criollo and Guianna populations. We also used the effective population size for *Medicago truncatula* from Siol et al. 2007 and the estimate of ρ from Paape et al. 2012, to calculate *r* for *M. truncatula* (=433 cM/Mb) and found that it was comparable with the mean rate found for the Criollo population (Table 1). One possible explanation for the higher recombination rate observed for the Criollo population is domestication; which has been observed to increase recombination rates, particularly in plants (Ross-Ibarra 2004). Previous work has shown that Criollo is the only population showing a strong signature of domestication, as revealed by its much higher drift parameter compared to other populations (Cornejo et al. 2018). The high recombination rate observed in Guianna (46.5 cM/Mb) can be explained in a similar way; while Guianna does not show a strong signature of domestication, it is the most recently established population (Bartley et al. 2005) and has undergone a recent bottleneck (Cornejo et al. 2018). We hypothesize from this result that the Guianna population is undergoing the initial stages of domestication, and its increased recombination is an early indicator of this. Another possibility is that the high recombination rates estimated for Criollo and Guianna can be explained by biases in estimation caused by errors associated to small samples or low genetic variation. However, the recombination rates for Amelonado (another population with low variation) and Purus (a population with small sample size) did not present this problem.

Analyses exploring mutations of putative recombination suppression genes (Fernandes et al. 2018b) could help disentangle the nature of this extreme variation in recombination rate in the Criollo and Guianna populations. It has been widely discussed that although actual recombination rates (chiasmata per bivalent) might increase in domestic plant populations, effective recombination is likely to be reduced in domestic populations as a result of increased LD and long runs of homozygosity. The interpretation for this observation is that it is likely that chromosomes might physically recombine at higher rates, but if homologous chromosomes contain the same sequence then no appreciable exchange of variants among different chromatid backgrounds can occur (Moyers et al. 2018). We find significantly higher estimates of median recombination rates for domesticated Criollo populations across the genome. Given previous research suggesting the key function of FIGL-1 on suppressing recombination rates in *Arabidopsis*, we investigated if mutations in this protein could explain the differences in recombination rate. Our results suggest that missense mutations in FIGL-1 could impair the activity of this protein in domesticated populations, reducing its efficacy as a recombination suppressor. The mutation showing the largest effect (KK_215) is close to the domain that has been previously shown to be involved in the interaction of FIGL-1 with RAD51 and DMC1 (Fernandes et al. 2018a) and thus lead us to propose that FIGL-1 might be responsible for the increase recombination rate in domesticated cacao populations. Further work using yeast two-hybrid systems similar to the work performed by Fernandes et al. (2018a) will be necessary in the future to show if the mutations do indeed interfere with the normal function of FIGL-1. These mutations were not found in increased frequency in Guianna, suggesting that the underlying mechanisms leading to increased recombination rates in this population are different to those in the Criollo population.

Despite recombination rates for eight of the ten populations being of the same order of magnitude, pairwise comparisons of average rates indicated that most populations have a significantly different rate of recombination from the others. The only exception were Nacional and Nanay whose average rates were not significantly different from each other. These two populations, however, are not more closely related to each other than they are to other populations, based on genetic differentiation (Cornejo et al. 2017). We interpret this result as suggestive that their similarity is not due to genetic similarity, but some other factors, e.g. epigenetics.

The likelihood of detecting hotspots of recombination in the genome will likely be affected by the amount of uncertainty in the estimates of recombination across sites or regions. Yet, we have been unable to identify any study where the magnitude of the uncertainty in the estimates of recombination are assessed to address this issue. We have performed careful comparisons and assessed the magnitude of the uncertainty in the estimation of recombination rates to show that this uncertainty is several orders of magnitude smaller than the variation in recombination rates across the genome (Table S1).

### Comparing recombination hotspot locations between populations

Understanding the pattern and rate of change of recombination hotspots at the population level can elucidate their role in shaping genome architecture, impacting how effectively selection operates (Felsenstein 1974). We found that a large proportion (55.5%) of hotspots detected are unique to a single population. The observed variability of hotspot location between populations points to demographic history not being the main driver of recombination hotspot location. However, the hotspots tend to appear in similar regions, as demonstrated by the Fisher’s exact tests (Table 2). This dichotomy can be explained by considering that the proportion of the genome occupied by recombination hotspots is very low, so even a small proportion of hotspots from two different populations being in the same region is enough for the Fisher’s exact test to recognize them as significantly similar. This small but significant similarity can occur by recombination being limited in its possible positioning along the genome, but not to the point of forcing hotspots to occur consistently in the same locations, and thus maintaining some level of stochasticity.

Given the significant proportion of overlapping hotspots between populations, it was still important to explore whether the similarities can be explained by shared genetic history. If demographic history explains the evolution of hotspot location, we would expect that more closely related populations would have a higher percent of overlapped hotspots. A significant relationship was found between population differentiation (F_ST_) and the distance between populations based on shared hotspots (Mantel test, *r=0.66*, *p=0.002*). This result indicates that, to some extent, the genetic differentiation and the location of hotspots are mirroring each other, which could be due to recombination hotspots being a product of the shared history between the populations. However, since recombination rates were estimated using a coalescent-based method, we expect historical relationships to be represented in our findings to some extent. We transformed the information of hotspot overlap to model hotspots as quantitative traits changing along a population tree (Cornejo et al. 2018). Our results show that a Brownian motion model (AIC=79.4) better fits the shared hotspot data than a model with stabilizing selection Ornstein-Uhlenbeck model (AIC=81.4), suggesting that, in principle, drift alone could explain the evolution of the location of recombination hotspots. However, the absolute number of hotspots that are shared among populations indicates that demographic history is insufficient to explain the evolution of recombination hotspots in this species.

While we do not detect all the hotspots in these populations and not all the hotspots detected are necessarily true positives, this proportion of unique hotspots can be seen as an indicator that the turnover rate for hotspots is faster than the time it took the 10 populations to differentiate. The detection rate for LDhot is approximately 55% under constant population conditions, and greater when a recent bottleneck has occurred (Auton et al. 2014, Dapper and Payseur 2017). Only two of the populations in this study (Criollo and Guianna) have a known recent bottleneck (Cornejo et al. 2018). However, Criollo is the only one of these two with an unusually low hotspot count (Table S4). Criollo’s low number of detected hotspots can be a product of its increased genome-wide recombination rate, making the signal of hotspots less pronounced. In a similar fashion, the increased overall recombination rate of Guianna may be affecting the detectability of hotspot regions, limiting our ability to resolve the limits of hotspots. It is important to note that our hotspots are unusually large for all populations (Table S4), which is likely a product of our low sample size leading to low resolution when resolving hotspot regions. Further work, increasing the sample size per population will contribute to increasing the resolution of these estimates.

While shared recombination hotspots can, to some extent, be explained by patterns of genetic differentiation, some of the sharing can simply be due to a tendency for hotspots to arise near TSSs and TTSs. It has been observed in other organisms that hotspots of recombination are frequently associated to specific genomic features (including TSSs and TTSs) (Auton et al. 2013, Choi et al. 2013, Hellsten et al. 2013, Myers et al. 2005, Singhal et al. 2015) or DNA sequence motifs (Auton et al. 2012, Brunschwig et al. 2012, Stevison et al. 2016). These factors can affect the landscape of recombination, contributing to the patterns of shared hotspot locations between populations that we observe in *T. cacao*.

Our ability to measure recombination in ten distinct populations allows us to analyze the relationship between population genetic processes and recombination. Our results suggest that the pattern of gains and losses of recombination hotspots is very dynamic and the landscape of recombination changes rapidly during the process of diversification within a species. This dynamism can have a tremendous impact on the adaptive dynamics of a species, and it should be taken into account, considering that theoretical studies tend to assume that recombination rates are constant during the evolution of populations (Hudson and Kaplan 1988, Donnelly and Kurtz 1999).

### Identifying DNA sequence motifs associated with the locations of recombination hotspots

The analysis of 17 hotspots shared between at least eight populations of *T. cacao* found an underrepresentation of retroelements and a marginal overrepresentation of DNA transposons when compared to the entire genome (Table 3). These results are not entirely surprising as it has already been suggested that transposable elements (TEs) tend to be enriched in areas of low recombination in *Drosophila* as a consequence of selection against TE activity that could lead to chromosome instability (Rizzon et al. 2002). However, the marginal over-representation of DNA transposons in the most conserved recombination hostspot is unexpected, given that all previous observations have shown a reduced representation of mobile elements in areas with high recombination rate (Rizzon et al. 2002). It is possible that DNA transposons are at least partly responsible for the maintenance of recombination hotspots as populations diverge, from which we expect that site-directed recombination is more frequent in these locations of the genome. However, the low percentage of these sequences observed in the set of all hotspots (Table 3) indicates that these sequences only have a small effect on the maintenance of hotspots. It has been observed in humans that short DNA motifs enriched for repeat sequences determine the location of 40 per cent of hotspots enriched for recurrent non-allelic homologous recombination (McVean 2010). One potential explanation for why natural selection does not eliminate hotspots in these regions is the possibility that these regions do not produce a large enough mutational load for natural selection to remove them from the population (McVean 2010 et al.).

### Identifying genomic features associated with the location of recombination hotspots

For all ten populations, an overrepresentation of hotspots was found in the areas immediately preceding and following transcribed regions of the chromosome. This matches the findings of previous studies in *Arabidopsis thaliana* (Choi et al. 2013), *Taenipygia guttata* and *Poephila acuticauda* (Singhal et al. 2015), and humans (Myers et al. 2005). The most likely explanation is that recombination events within genes are selected against. The rationale being that a recombinant chromosome that undergoes a double-strand break in the middle of a coding region will have a higher risk of being inviable, and therefore not represented in the current set of chromosomes for its population. Recombination occurring in transcription start and stop sites, on the other hand, does a much better job at breaking up haplotypes or shuffling alleles in different genomic backgrounds, while preserving the functionality of coding regions. This rationale is supported by previous findings of increased recombination rates in these regions (Choi et al. 2013). It is also supported by results from PRDM9 knock-out *Mus musculus*, which has shown a reversion to hotspots located near TSSs (Brick et al. 2011). The enrichment of *T. cacao* hotspots in TSSs and TTSs is thus a reasonable result given that zinc-finger binding motifs and potential modifiers like PRDM9 have not been identified in this species.

### Implications for the evolutionary history of T. cacao

Overall, our results show a large, consistent pattern where recombination rates in the ten populations of *T. cacao* are of a similar magnitude as mutation rates but show a high diversity in location and number of hotspots of recombination that cannot be explained solely by the process of diversification of the populations. A potential hypothesis that could explain the rapid turnover of hotspots of recombination and the relative differences in recombination among populations is that epigenetic changes are involved in controlling the turnover of recombination in plants. This hypothesis is not unreasonable given the recent observation of epigenetic control of recombination in plants (Yelina et al. 2015). Further theoretical and simulation work should be done in order to better understand the implications of the rapidly changing recombination hotspots in adaptive dynamics. We also show that there is an overall underrepresentation of hotspots in exons and introns for most populations, which is consistent with purifying selection acting against changes that could result in disruptions of gene function. On the other hand, we observed an overrepresentation of hotspots in TTSs and TSSs for all ten populations. This could impact the maintenance and spread of beneficial traits in the population by shuffling allelic variants of genes without causing disruption of their function. We hypothesize that the enrichment of hotspots of recombination in TTSs and TSSs can have an important impact in the spread of beneficial mutations across different genomic backgrounds; increasing the rate of adaptation to selective pressures (e.g. selection for improved pathogen response).

## Materials and Methods

### Comparing recombination rates between populations

Sequence data were downloaded from the Cacao Genome Database and NCBI (Accession PRJNA486011), including the reference sequence for each chromosome and the full genome annotation (*Theobroma cacao* cv. Matina 1-6 v1.1) (Motamayor et al. 2013). Processing was done using the pipeline from (Cornejo et al 2018) available at the github repository *oeco28/Cacao_Genomics*. Full genome data was used from a total of 73 individuals (146 chromosomes) across 10 populations (Cornejo et al. 2018). We filtered single nucleotide polymorphism data and excluded rare variants (minor allele frequency <= 0.05) per population. Separate variant files per population per chromosome were then phased using default conditions with SHAPEIT2 (Delaneau et al. 2011) under default parameters. Haplotype files were converted back to phased variant calling format (vcf) for its downstream analysis. We have also phased the data with Beagle (Browning and Browning. 2007), using a burn-in of 10000 iterations, and estimations done over 10000 iterations. No appreciable differences were observed between the two methods and Beagle phasing was maintained for the analyses. The reason for performing the phasing separately for each population is that linkage disequilibrium patterns are expected to be affected by population structure. The ten populations have been shown to be unique clusters with very little admixture between them (Cornejo et al. 2018), and the individuals used in this study were those whose ancestry was clearly from a single population. VCFTools (Danecek et al. 2011) was used to remove all singletons and doubletons (i.e. SNPs where the minor allele count is less than three). Only bi-allelic single nucleotide polymorphisms (SNPs) were retained and were exported in LDhat format. The sample size and post-filtering SNP count for each population can be found in Table S5.

In order to estimate recombination rates, we used the *interval* routine from LDhat (Auton and McVean 2007), a program that implements coalescent resampling methods to estimate historical recombination rates from SNP data. To reduce computation time, each chromosome was split into windows, each containing 2000 SNPs. To counteract the overestimation of recombination rate produced at the ends of the windows, an overlap of 500 SNPs was left between consecutive windows. The final window for each chromosome did not always match the general scheme, so the final 2000 SNPs were taken (making the overlap with the second to last window variable, but never less than 500 SNPs) (Fig. S4). Once these windows were generated, LDhat was run over each window using 100 million iterations, sampling every 10000 iterations (10000 total points sampled), with a block penalty of 5. Lookup tables with a grid of 100 points, population mutation rate parameters (θ) of 0.1 and 0.001, and a number of sequences (n) of 50 were used for all populations. Downstream analyses were conducted using the results from the θ=0.1 runs of LDhat. We used the same θ for all populations since estimates from (Cornejo et al. 2018) ranged from π=0.27% to π=0.37%, all comfortably within an order of magnitude of each other. The first 50 million iterations were discarded as burn-in. Once recombination rates were calculated, 250 positions were cut off from both windows involved in each overlap, so that the estimates for the first half of the overlap was taken from the end of the preceding window and the estimates for the second half of the overlap were taken from the beginning of the following window. The final overlap in each chromosome was split in order to remove 250 SNPs from the second to last window, regardless of the remaining size of the last window. The remaining rate estimates were then merged in order to obtain recombination rates for the entire chromosome. This was done for each chromosome of each population.

The estimation of recombination rates with LDhat is approximated using a sampling scheme with a Markov Chain Monte Carlo (MCMC) algorithm as implemented in the *interval* routine. The inference of recombination rates is the result of the integration of estimated parameter values across iterations with the routine *stats*. In the majority of recent studies where LDhat or LDhelmet are used (Myers et al. 2005, Auton et al. 2012, Brunschwig et al. 2012, Paape et al. 2012, Auton et al. 2013, Choi et al. 2013, Singhal et al. 2015, Stevison et al. 2016), whether there is convergence of the Markov chains has not been explicitly investigated. One study that we are aware of has used simulations to assess whether their small sample size affected their ability to obtain reliable estimates of recombination using LDhelmet (Booker et al. 2017) but did not assess the uncertainty of the estimates from the MCMC process itself. We argue that evaluation of convergence is important to assess the confidence in the estimated reported values, especially if there is interest in analyzing the differences in recombination rate along the genome. Visual inspection of pilot runs of the analysis demonstrated that convergence was not achieved after running 40M iterations, which is why the length of the chains was increased to 100M iterations. Additionally, we explored the uncertainty in the estimates of recombination site-wise by integrating over the trace of the estimates for recombination rate to infer the 95% Credibility Interval. We then estimated the 95% interval of recombination estimates range across all sites in the genome to have an overall measure of uncertainty that we compared to the median 95% Credibility Interval for the trace of each position.

In order to compare recombination rates, the effective population size (*N*_*e*_) calculated for each population (Cornejo et al. 2018) was used to convert rates in *N*_*e*_*r* to *r*. We tested the difference between the whole-genome mean rate of recombination (*r*) between populations using the Wilcoxon signed-rank test `wilcox.test` function from the `stats` package in `R`) (R core team 2018). There were 45 comparisons, making the Bonferroni correction cutoff value: α=0.0011.

Suppressors of recombination identified in Arabidopsis and other systems, FIGL1 and FLIP, have not yet been identified in *T. cacao* (Motamayor et al. 2013). We performed a reciprocal BLAST search using a newly generated databased containing sequences of FIGL1 and FLIP obtained from ncbi (Altschul et al. 1990, Altschul et al. 1994b, Shiryev et al. 2007). After identification of orthologous copies of FIGL1 and FLIP, we extracted the annotated variants responsible for missense mutations in the proteins from data generated in Cornejo et al. 2018. We then used these mutations to estimate the frequency of homozygous genotypes with alternative alleles (different to the reference). Because the reference genome belongs to the Amelonado populations, the population that presents the lowest estimated median recombination rate in our work, we used the frequency of homozygous alternatives to infer the impact of missense mutations under the assumption of a recessive model on the recombination rate. For this, we estimated the homozygosity for missense mutations and eliminated those found in complete correlation (r^2^ = 1) and fitted a generalized linear model of the form:

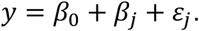

Where *y* is a vector of the median recombination rate accross populations, b_0_ is the intersect (in this case the Amelonado median recombination rate) and b_j_ is the effect of the homozygosity in position *j*. We assumed a Gaussian link function for the model.

### Comparing recombination hotspot locations between populations

Recombination hotspots were estimated with LDhot (Auton and McVean 2007), a likelihood-based program that tests whether a single distribution model or a two-distribution model better explains the observed recombination rates in 1 kb sliding windows (default), for each chromosome. Each chromosome was run in its entirety, with the number of simulations (nsims) set to 1000. The resulting potential hotspots were refined by an α of 0.001, and overlapping hotspots were merged. This method therefore detects hotspots by comparing rates in 1 kb windows to the rates in the surrounding regions.

To determine the set of consensus hotspots, the hotspots from all populations were merged. Two hotspots from different populations were considered to be shared if they both overlapped with the same hotspot in the consensus set. To summarize all shared hotspots, a Boolean matrix was constructed, in which a population having a hotspot that overlaps with a hotspot in the consensus list leads to an indication of presence of the consensus hotspot in that population. This matrix was used to determine hotspots shared by two or more populations.

A Fisher’s exact test was run for each pair of populations in order to determine whether hotspots for the pair of populations overlap significantly more than expected. The BED files containing the location of the recombination hotspots for each pair of populations were compared using Bedtools:fisher (Quinlan and Hall. 2010). The number of comparisons was 45, making the Bonferroni correction cutoff value: α=0.0011.

In order to compare the relationships between populations based on shared hotspots we calculated Jaccard distances (`distance` function, `philentropy` package, `R`) (Drost 2018) and compared them to a published F_ST_ matrix (Cornejo et al. 2018) using a Mantel test (`mantel.rtest` function, `ade4` package, `R`) (Chessel et al. 2004, Dray et al. 2007a, Dray et al. 2007b, Bougeard and Dray 2018). The F_ST_ estimates from (Cornejo et al. 2018) were generated using Weir and Cockerham’s estimator (Weir and Cockerham 1984).

In order to model the presence or absence of hotspots along a drift tree, a multiple correspondence analysis was used on the Boolean matrix of shared hotspots using the `MCA` function from the `FactoMineR` package in `R` (Le et al. 2008). Nine dimensions were retained and used as traits along a previously generated drift tree (Cornejo et al. 2018). Using the `Rphylopars` package in `R` (Goolsby et al. 2016), the dimensions were modeled as Brownian motion and as an Ornstein-Uhlenbeck process. The fit of the two models were compared using the AIC values for the best fitting models of each type.

### Identifying DNA sequence motifs associated with the locations of recombination hotspots

Motifs associated with hotspots were found using RepeatMasker (Smith et al. 2016). The entire genome, the set of consensus hotspots, and a set of ubiquitous hotspots (hotspots shared by at least eight of the populations) were examined with RepeatMasker, using normal speed and "theobroma cacao" in the species option. In order to determine whether ubiquitous hotspots were enriched for particular DNA sequences, a set of the same number and size of sequences was randomly selected from the genome using Bedtools:shuffle (Quinlan and Hall. 2010) and examined with RepeatMasker. This simulation was repeated one thousand times and a null distribution against which observed values were compared was constructed from the results.

### Identifying genomic features associated with the location of recombination hotspots

Testing whether recombination hotspots were overrepresented near particular genomic features was done by using a resampling scheme to establish null expectations and then comparing the observed value to the empirical distribution. For each feature, locations were retrieved and the number of observed hotspots that overlap with this feature were counted. To determine whether this number of overlapping hotspots was unusually high or low, a set of hotspots that matched the number of hotspots and the size of each hotspot was simulated. These simulated hotspots were placed randomly along the chromosome, using a uniform distribution. The simulation was run 1000 times and the number of simulated hotspots that overlap with the true genomic features was measured for each simulation. The simulations generate an expected distribution of overlap with the genomic feature, and the true value was then compared to the distribution. When simulated hotspots overlapped, the location of one of them was sampled again. Features tested were: Transcriptional start sites (TSSs), transcriptional termination sites (TTSs), exons, and introns. TSSs and TTSs are considered to be the 500bp upstream and downstream of coding regions respectively.

The reason for the proposed novel resampling scheme is that, if the size and distribution of genomic features and hotspots were not taken into account, it would set unrealistic expectations for the overlap between features under a null model of no association. In this sense, the null model would be inappropriate and potentially inflate the false positive rate.

## Data Access

Rate and summary files from LDhat runs as well as hotspots for each population will be placed in a Dryad repository. Scripts for LDhat and LDhot runs as well as the resampling schemes used and additional analysis is available in the following github repository: *ejschwarzkopf/recombination-map*.

## Acknowledgements

The authors would like to thank the Noe Higinbotham endowment and the WSU College of Arts and Science for travel funds to EJS to present earlier versions of this work. We would like to thank the Kamiak High Performance Computing Cluster at WSU for the infrastructure support to run the analyses, and the Cornejo, Kelley, and Busch labs at WSU for feedback and edits on the manuscript.

## Supplementary Materials

**Table S1.** The median of the upper and lower bounds of the 95% Credibility Interval for the trace of estimates of *r* (in cM/Mb) from all positions in the genome are presented for each population (i.e. Position L95 and Position U95). The upper and lower bounds of the 95% probability interval for the median estimate of *r* for each population is also presented (i.e. Genome L95 and Genome U95). The quotients of the upper and lower bounds for each of the two intervals point to a much larger genome-wide variation in *r* than per-position variation in the trace for the estimate of *r*.

| Population | Position L95 | Position U95 | Genome L95 | Genome U95 | Position Range Quotient | Genome Range Quotient |
| --- | --- | --- | --- | --- | --- | --- |
| Amelonado | 6.35e-05 | 7.67e-03 | 2.33e-05 | 31.3 | 120.75 | 1.34e+06 |
| Contamana | 1.63e-01 | 1.34 | 1.40e-04 | 26.4 | 8.22 | 1.88e+05 |
| Criollo | 87.5 | 43.1 | 5.35e-02 | 1,660 | 4.92 | 3.11e+04 |
| Curaray | 5.15e-01 | 2.63 | 2.72e-04 | 20.2 | 5.11 | 7.40e+04 |
| Guianna | 9.76e-01 | 12.6 | 1.81e-04 | 296 | 12.90 | 1.63e+06 |
| Iquitos | 5.50e-05 | 6.52e-03 | 1.58e-05 | 24.5 | 186.29 | 1.55e+06 |
| Maranon | 1.98e-04 | 7.40e-02 | 2.31e-05 | 35.2 | 373.35 | 1.52e+06 |
| Nacional | 4.80e-05 | 6.50e-03 | 2.76e-05 | 36.6 | 135.60 | 1.33e+06 |
| Nanay | 1.65e-04 | 3.06e-02 | 2.06e-05 | 35.2 | 185.32 | 1.71e+06 |
| Purus | 3.98e-03 | 5.24e-01 | 2.00e-04 | 63.5 | 131.87 | 3.18e+05 |

**Table S2.**
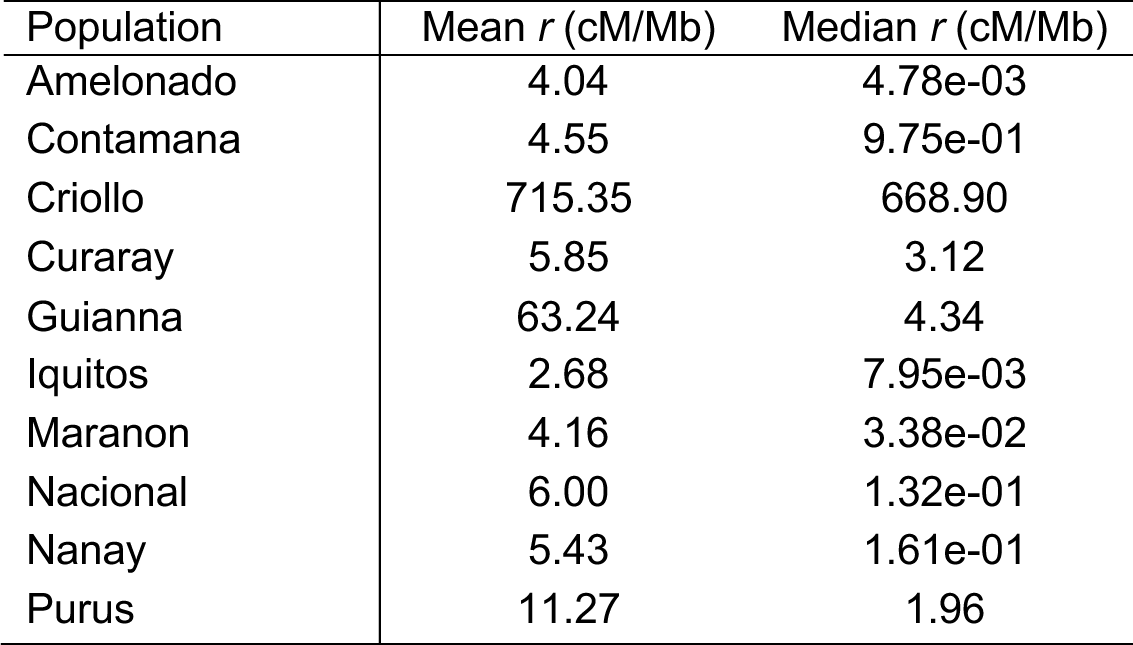
Mean and median genome-wide recombination rates (*r*) in cM/Mb for all ten *T. cacao* populations obtained using LDhat with θ= 0.001.

**Table S3.** Name of *T. cacao* gene coding for FIGL1 and FLIP and amino acid mutations for FIGL1 and FLIP orthologs.

| Protein | FIGL1 | FLIP |
| --- | --- | --- |
| Gene | *Thecc1EG017182t2* | *Thecc1EG020410t1* |
| Mutations | K9Q, F66L, D114Y, T155I, D189V, K195M, E215K, S235N, R270Q, A285S, S291T, V548I, G566S, A567V, M580T | L102F, S153T, K174T, Y140N, V125A, H203R, Q237K, V406L, S430Y, R446Q, M497I, L521M, D696V, L729I, M739I, A746G, K875Q, P906L |

**Table S4.** Average hotspot size (in kb) and count for hotspots detected in each population and average for all populations.

| Population | Mean hotspot size (kb) | Hotspot count |
| --- | --- | --- |
| Amelonado | 6.9 | 1325 |
| Contamana | 6.1 | 5184 |
| Criollo | 6.1 | 887 |
| Curaray | 5.8 | 2303 |
| Guianna | 8.6 | 3655 |
| Iquitos | 7.0 | 3258 |
| Maranon | 6.8 | 3296 |
| Nacional | 6.9 | 2202 |
| Nanay | 7.6 | 3818 |
| Purus | 6.3 | 3972 |
| Average | 6.9 | 2989.9 |

**Table S5.** Sample size and post-filtering SNP count for all ten populations of *Theobroma cacao* for which recombination maps were generated. The proportion of the genome that is callable is also reported.

| Population | Sample size (chromosome) | SNP count | Proportion callable |
| --- | --- | --- | --- |
| Amelonado | 11 (22) | 373,789 | 0.68 |
| Contamana | 9 (18) | 2,097,618 | 0.87 |
| Criollo | 4 (8) | 309,818 | 0.69 |
| Curaray | 5 (10) | 1,106,871 | 0.70 |
| Guianna | 9 (18) | 770,729 | 0.915 |
| Iquitos | 7 (14) | 1,575,711 | 0.922 |
| Maranon | 14 (28) | 1,783,226 | 0.955 |
| Nacional | 4 (8) | 718,099 | 0.89 |
| Nanay | 10 (20) | 830,885 | 0.94 |
| Purus | 6 (12) | 1,184,181 | 0.9 |

**Figure S1.**
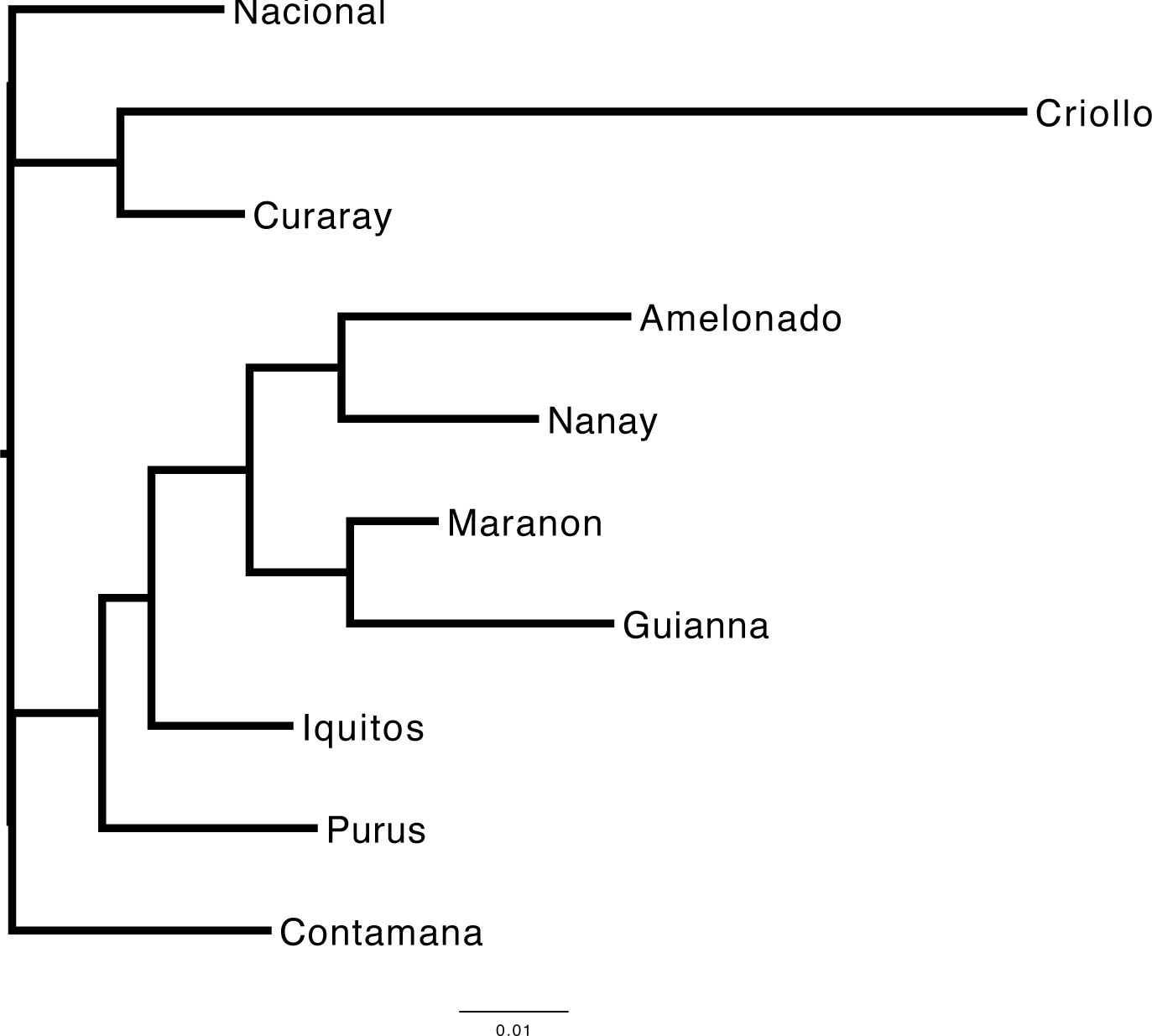
Drift tree constructed using treemix (Pickrell and Pritchard, 2012) for the 10 *T. Cacao* populations. Distances between populations are based on the drift parameter. Modified from Cornejo et al. (2018)

**Figure 2.**
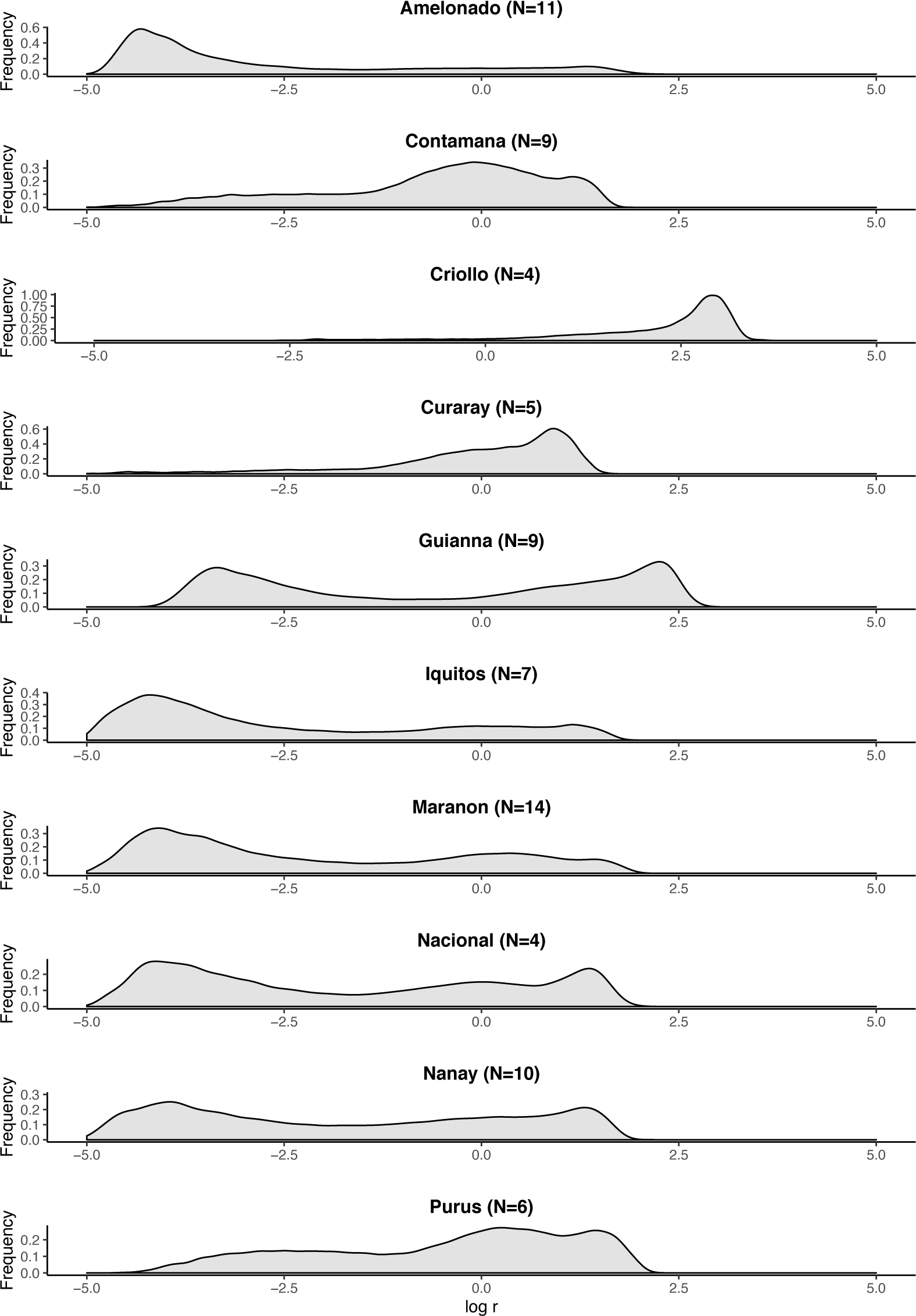
Distribution of *log*_*10*_ recombination rates (*log*_*10*_*(r)*) along the genomes of the ten *T. Cacao* populations. The sample size (N) is reported for each population.

**Figure S3.**
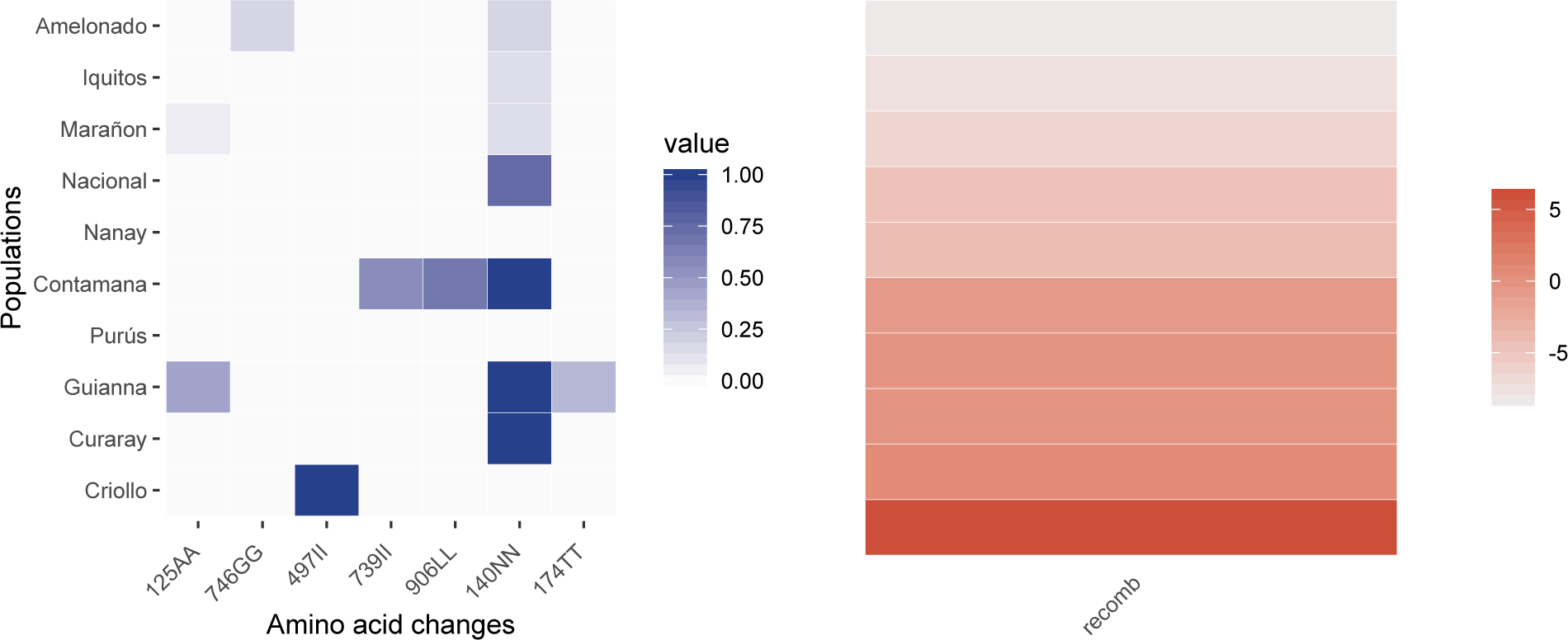
The left panel shows the frequency of individuals that are homozygous for the alternative allele of amino acid mutations in a *T. cacao* FLIP ortholog. Alternative allele is defined in terms of the Amelonado reference genome. The right panel shows the *log*_*e*_ transformed recombination rates (*r*). The populations are in the same order in both panels.

**Figure S4.**
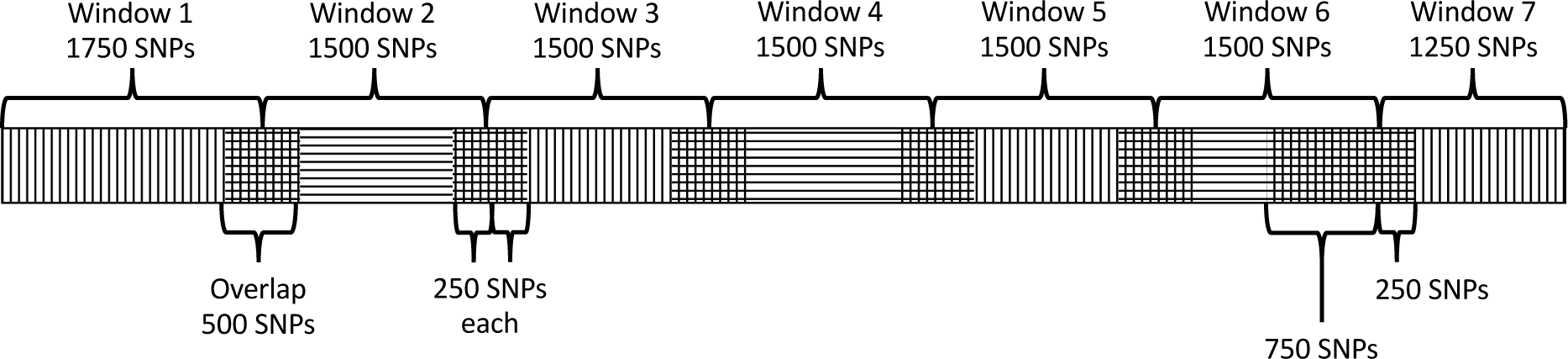
Example of the window layout for a 10,750 SNP chromosome. The 2,000 SNP long windows are represented by alternating horizontal and vertical lines and the overlaps between them are represented by square crosshatches. Braces above the chromosome indicate the regions from which recombination rates are extracted to generate the chromosome-wide recombination rates.

## References

Akhunov E, Goodyear A, Geng S, Qi L, Echalier B, Gill B, Gustafson J, Lazo G, Chao S, Anderson O, et al. 2003. The Organization and Rate of Evolution of Wheat Genomes Are Correlated With Recombination Rates Along Chromosome Arms. Genome Res. 13:753–763. Doi:10.1101/gr.808603.

Altschul S, Gish W, Miller W, Myers E, Lipman D. 1990. Basic local alignment search tool. J Mol Biol. 215:403–410.

Altschul S, Boguski M, Gish W, Wootton J. 1994. Issues in searching molecular sequence databases. Nature Genet. 6:119–129

Anderson L, Salameh N, Bass H, Harper L, Cande W, Weber G, Stack S. 2004. Integrating Genetic Linkage Maps With Pachytene Chromosome Structure in Maize. Genetics. 166:1923–1933.

Auton A, McVean G. 2007. Recombination rate estimation in the presence of hotspots. Genome res. 17:1219–1227.

Auton A, Myers S, McVean G. 2014. Identifying recombination hotspots using population genetic data. bioRxiv Doi:1403.4264v1.

Auton A, Fledel-Alon A, Pfeifer S, Venn O, Ségurel L, Street T, Leffler EM, Bowden R, Aneas I, Broxholme J, et al. 2012. A fine-scale chimpanzee genetic map from population sequencing. Science. 336:193–198. Doi:10.1126/science.1216872.

Auton A, Rui Li Y, Kidd J, Oliveira K, Nadel J, Holloway JK, Hayward JJ, Cohen PE, Greally JM, Wang J, et al. 2013. Genetic Recombination Is Targeted towards Gene Promoter Regions in Dogs. PLoS Genet. 9. Doi:10.1371/journal.pgen.1003984.

Bartley BGD. 2005. The genetic diversity of cacao and its utilization. CABI. Wallingford, United Kingdom.

Begun DJ, Aquadro CF. 1992. Levels of naturally occurring DNA polymorphism correlate with recombination rates in *D. melanogaster*. Nature. 356. Doi:10.1038/356519a0.

Booker TR, Ness RW, Keightley PD. 2017. The Recombination Landscape in Wild House Mice Inferred Using Population Genomic Data. Genetics. 207:297–309. Doi:10.1534/genetics.117.300063.

Bougeard S, Dray S. 2018. Supervised Multiblock Analysis in R with the ade4 Package. J Stat Softw. 86:1–17. Doi:10.18637/jss.v086.i01.

Branca A, Paape T, Zhou P, Briskine R, Farmer A, Mudge J, Bharti A, Woodward J, May G, Gentzbittel L, et al. 2011. Whole-genome nucleotide diversity, recombination, and linkage disequilibrium in the model legume Medicago truncatula. P Natl Acad Sci USA. 108:E864–E870.

Brick K, Smagulova F, Khil P, Camerini-Otero RD, Petukhova GV. 2012. Genetic recombination is directed away from functional genomic elements in mice. Nature. 485:642–645. Doi:10.1038/nature11089.

Browning SR, Browning BL. 2007. Rapid and accurate haplotype phasing and missing-data inference for whole-genome association studies by use of localized haplotype clustering. Am J Hum Genet. 81.

Brunschwig H, Levi L, Ben-David E, Williams R, Yakir B, Shifman S. 2012. Fine-Scale Maps of Recombination Rates and Hotspots in the Mouse Genome. Genetics. 191:757–64.

Chessel D, Dufour A, Thioulouse J. 2004. The ade4 Package – I: One-Table Methods. R News, 4, 5–10.

Choi K, Zhao X, Kelly K, Venn O, Higgins J, Yelina N, Hardcastle T, Ziolkowski P, Copenhaver G, Franklin F, et al. 2013. *Arabidopsis* meiotic crossover hot spots overlap with H2A.Z nucleosomes at gene promoters. Nature Genet. 45:1327–36.

Cornejo OE, Yee M-C, Dominguez V, Andrews M, Sockell A, Strandberg E, Livingstone D, Stack C, Romero A, Umaharan P, et al. 2018. Population genomic analyses of the chocolate tree, *Theobroma cacao* L., provide insights into its domestication process. Commun biol. 1:167–167. Doi:10.1038/s42003-018-0168-6.

Crow JF, Kimura, Motoo. 1970. An introduction to population genetics theory. Harper & Row, New York.

Danecek P, Auton A, Abecasis G, Albers CA, Banks E, DePristo MA, Handsaker RE, Lunter G, Marth GT, Sherry ST, et al. 2011. The variant call format and VCFtools. Bioinformatics. 27:2156–2158. Doi:10.1093/bioinformatics/btr330.

Dapper AL, Payseur BA. 2018. Effects of Demographic History on the Detection of Recombination Hotspots from Linkage Disequilibrium. Mol Biol Evol. 35:335–353. Doi:10.1093/molbev/msx272.

Delaneau O, Marchini J, Zagury J. 2012. A linear complexity phasing method for thousands of genomes. Nat Methods. 9:179–81. Doi:10.1038/nmeth.1785.

Donnelly P, Kurtz TG. 1999. Genealogical Processes for Fleming-Viot Models with Selection and Recombination. Ann Appl Probab. 9:1091–1148.

Dray S, Dufour A. 2007. The ade4 Package: Implementing the Duality Diagram for Ecologists. J Stat Softw, 22:1–20. Doi:10.18637/jss.v022.i04.

Dray S, Dufour A, Chessel D. 2007. The ade4 Package – II: Two-Table and K-Table Methods. R News, 7:47–52.

Drost HG. 2018. Philentropy: Information Theory and Distance Quantification with R. J Open Source Softw. 3:765.

Eyre-Walker A, Keightley P. 2007. The distribution of fitness effects of new mutations. Nat Rev Genet. 8:610–618. Doi:10.1038/nrg2146.

Felsenstein J. 1974. The evolutionary advantage of recombination. Genetics. 78:737–56.

Fernandes J, Duhamel M, Seguéla-Arnaud M, Froger N, Girard C, Choinard S, et al. 2018a. FIGL1 and its novel partner FLIP form a conserved complex that regulates homologous recombination. PLoS Genet. 14:e1007317. Doi:10.1371/journal.pgen.1007317

Fernandes JB, Séguéla-Arnaud M, Larchevêque C, Lloyd AH, Mercier R. 2018b. Unleashing meiotic crossovers in hybrid plants. P Natl Acad Sci USA. 115:2431–2436. Doi:10.1073/pnas.1713078114.

Girard C, Chelysheva L, Choinard S, Froger N, Macaisne N, Lehmemdi A, et al. 2015. AAA-ATPase FIDGETIN-LIKE 1 and Helicase FANCM Antagonize Meiotic Crossovers by Distinct Mechanisms. PLoS Genet. 11: e1005369. Doi:10.1371/journal.pgen.1005369

Goolsby EW, Bruggeman J, Ané C. 2017. Rphylopars: fast multivariate phylogenetic comparative methods for missing data and within-species variation. Methods in Ecol Evol. 8:22–27. Doi:10.1111/2041-210X.12612.

Gore MA, Chia J-M, Elshire RJ, Sun Q, Ersoz ES, Hurwitz BL, Peiffer JA, Mcmullen MD, Grills GS, Ross-Ibarra J, et al. 2009. A First-Generation Haplotype Map of Maize. Science. 326:1115–1117. Doi:10.1126/science.1177837.

Haldane J. 1937. The Effect of Variation on Fitness. Am Nat. 71.

Hellsten U, Wright K, Jenkins J, Shu S, Yuan Y, Wessler S, Schmutz J, Willis J, Rokhsar D. 2013. Fine-scale variation in meiotic recombination in *Mimulus* inferred from population shotgun sequencing. P Natl Acad Sci USA. 110.

Henderson J, Joyce R, Hall G, Hurst W, McGovern P. 2007. Chemical and archaeological evidence for the earliest cacao beverages. P Natl Acad Sci USA. 104.

Hinch A, Tandon A, Patterson N, Song Y, Rohland N, Palmer C, Chen G, Wang K, Buxbaum S, Akylbekova E, et al. 2011. The landscape of recombination in African Americans. Nature. 476:170–5. Doi:10.1038/nature10336.

Hudson RR, Kaplan NL. 1988. The coalescent process in models with selection and recombination. Genetics. 120:831–840.

Kim S, Plagnol V, Hu T, Toomajian C, Clark R, Ossowski S, Ecker J, Weigel D, Nordborg M. 2007. Recombination and linkage disequilibrium in *Arabidopsis thaliana*. Nat Genet. 39:1151–1155. Doi:10.1038/ng2115.

Lê S, Josse J, Husson F. 2008. FactoMineR: An R Package for Multivariate Analysis. J Stat Softw. 25. Doi:10.18637/jss.v025.i01.

Li W-H, Nei M. 1974. Stable linkage disequilibrium without epistasis in subdivided populations. Theor Popul Biol. 6:173–183. Doi:10.1016/0040-5809(74)90022-7.

Luo C, Shu B, Yao Q, Wu H, Xu W, Wang S. 2016. Construction of a High-Density Genetic Map Based on Large-Scale Marker Development in Mango Using Specific-Locus Amplified Fragment Sequencing (SLAF-seq). Front Plant Sci. 7. Doi:10.3389/fpls.2016.01310.

Mackiewicz D, de Oliveira PMC, Moss de Oliveira S, Cebrat S, Lustig AJ. 2013. Distribution of Recombination Hotspots in the Human Genome – A Comparison of Computer Simulations with Real Data. PLoS ONE. 8. Doi:10.1371/journal.pone.0065272.

Mackiewicz D, Zawierta M, Waga W, Cebrat S. 2010. Genome analyses and modelling the relationships between coding density, recombination rate and chromosome length. J Theor Biol. 267:186–192. Doi:10.1016/j.jtbi.2010.08.022.

Martinez-Perez E, Schvarzstein M, Barroso C, Lightfoot J, Dernburg AF, Villeneuve AM. 2008. Crossovers trigger a remodeling of meiotic chromosome axis composition that is linked to two-step loss of sister chromatid cohesion. Gene Dev. 22:2886–901. Doi:10.1101/gad.1694108.

McVean G. 2010. What drives recombination hotspots to repeat DNA in humans? Philos T Roy Soc B. 365:1213–1218. Doi:10.1098/rstb.2009.0299.

Mcvean G, Myers S, Hunt S, Deloukas P, Bentley D, Donnelly P. 2004. The fine-scale structure of recombination rate variation in the human genome. Science. 304:581–4.

Mézard C. 2006. Meiotic recombination hotspots in plants. Biochem Soc T. 34:531–4.

Motamayor J, Lachenaud P, Loor R, Kuhn D, Brown J, Schnell R. 2008. Geographic and Genetic Population Differentiation of the Amazonian Chocolate Tree (*Theobroma cacao* L). PLoS One. 3. Doi:10.1371/journal.pone.0003311.

Motamayor J, Mockaitis K, Schmutz J, Haiminen N, Livingstone D, Cornejo O, Findley S, Zheng P, Utro F, Royaert S, et al. 2013. The genome sequence of the most widely cultivated cacao type and its use to identify candidate genes regulating pod color. Genome Biol. 14:r53–r53. Doi:10.1186/gb-2013-14-6-r53.

Mousavi M, Tong C, Liu F, Tao S, Wu J, Li H, Shi J. 2016. De novo SNP discovery and genetic linkage mapping in poplar using restriction site associated DNA and whole-genome sequencing technologies. BMC Genomics. 17. Doi:10.1186/s12864-016-3003-9.

Moyers B, Morrell P, McKay J. 2018. Genetic Costs of Domestication and Improvement. J Hered. 109: 103–116.

Myers S, Bottolo L, Freeman C, McVean G, Donnelly P. 2005. A fine-scale map of recombination rates and hotspots across the human genome. Science. 310:321–324.

Ohta T. 1982. Linkage disequilibrium due to random genetic drift in finite subdivided populations. P Natl Acad Sci USA. 79:1940–1944. Doi:10.1073/pnas.79.6.1940.

Ollitrault P, Terol J, Chen C, Federici C, Lotfy S, Hippolyte I, Ollitrault F, Bérard A, Chauveau A, Cuenca J, et al. 2012. A reference genetic map of *C. clementina* hort. ex Tan.; citrus evolution inferences from comparative mapping. BMC Genomics. 13. Doi:10.1186/1471-2164-13-593.

Otto S, Barton N. 1997. The evolution of recombination: removing the limits to natural selection. Genetics. 147:879–906.

Paape T, Zhou P, Branca A, Briskine R, Young N, Tiffin P. 2012. Fine-scale population recombination rates, hotspots, and correlates of recombination in the *Medicago truncatula* genome. Genome Biol Evol. 4:726–737. Doi:10.1093/gbe/evs046.

Pickrell J, Pritchard J. 2012. Inference of Population Splits and Mixtures from Genome-Wide Allele Frequency Data. PLoS Genetics. 8. doi:10.1371/journal.pgen.1002967.

Ptak S, Hinds D, Koehler K, Nickel B, Patil N, Ballinger D, Przeworski M, Frazer K, Pääbo S. 2005. Fine-scale recombination patterns differ between chimpanzees and humans. Nature Genet. 37:429–434.

Quinlan A, Hall I. 2010. BEDTools: a flexible suite of utilities for comparing genomic features. Bioinformatics. 26:841–842. Doi:10.1093/bioinformatics/btq033.

R Core Team. 2014. R: A language and environment for statistical computing. R Foundation for Statistical Computing, Vienna, Austria. URL http://www.R-project.org/.

Revelle W. 2019. psych: Procedures for Psychological, Psychometric, and Personality Research. Northwestern University, Evanston, Illinois. R package version 1.9.12, https://CRAN.R-project.org/package=psych.

Rizzon C, Marais G, Gouy M, Biémont C. 2002. Recombination rate and the distribution of transposable elements in the Drosophila melanogaster genome. Genome Res. 12:400–407.

Rodgers K, McVey M. 2016. Error-Prone Repair of DNA Double-Strand Breaks. J Cell Physiol. 231:15–24. Doi:10.1002/jcp.25053.

Ross-Ibarra J. 2004. The Evolution of Recombination under Domestication: A Test of Two Hypotheses. Am Nat. 163:105–112. Doi:10.1086/380606.

Sanjuán R, Moya A, Elena SF, Ohta T. 2004. The Distribution of Fitness Effects Caused by Single-Nucleotide Substitutions in an RNA Virus. P Natl Acad Sci USA. 101:8396–8401.

Schnable P, Ware D, Fulton R, Stein J, Wei F, Pasternak S, Liang C, Zhang J, Graves L, Minx T, et al. 2009. The B73 maize genome: complexity, diversity, and dynamics. Science. 326:1112–1115. Doi:10.1126/science.1178534.

Shanfelter A, Archambeault S, White M, Hurst L. 2019. Divergent Fine-Scale Recombination Landscapes between a Freshwater and Marine Population of Threespine Stickleback Fish. Genome Biol Evol. 11:1573–1585. Doi:10.1093/gbe/evz090.

Shiryev S, Papadopoulos J, Schäffer A, Agarwala R. 2007. Improved BLAST searches using longer words for protein seeding. Bioinformatics. 23:2949–51.

Singhal S, Leffler E, Sannareddy K, Turner I, Venn O, Hooper D, Strand A, Li Q, Raney B, Balakrishnan C, et al. 2015. Stable recombination hotspots in birds. Science. 350:928–932. Doi:10.1126/science.aad0843.

Siol M, Bonnin I, Olivieri I, Prosperi J, Ronfort J. 2007. Effective population size associated with self-fertilization: lessons from temporal changes in allele frequencies in the selfing annual Medicago truncatula. J Evolution Biol. 20:2349–2360. Doi:10.1111/j.1420-9101.2007.01409.x.

Smith A, Hubley R, Green P. 2016. RepeatMasker Open-4.0 (2013-2015). <http://www.repeatmasker.org>.

Stapley J, Feulner P, Johnston S, Santure A, Smadja C. 2017. Variation in recombination frequency and distribution across eukaryotes: patterns and processes. Philos T Roy Soc B. 372. Doi:10.1098/rstb.2016.0455.

Stevison L, Woerner A, Kidd J, Kelley J, Veeramah K, McManus K, Bustamante C, Hammer M, Wall J. 2016. The Time Scale of Recombination Rate Evolution in Great Apes. Mol Biol Evol. 33:928–945. Doi:10.1093/molbev/msv331.

Weir B, Cockerham C. 1984. Estimating f-statistics for the analysis of population structure. Evolution. 38:1358–1370. Doi:10.1111/j.1558-5646.1984.tb05657.x.

Winckler W, Myers S, Richter D, Onofrio R. 2005. Comparison of Fine-Scale Recombination Rates in Humans and Chimpanzees. Science. 308:107–11.

Wloch D, Szafraniec K, Borts R, Korona R. 2001. Direct estimate of the mutation rate and the distribution of fitness effects in the yeast Saccharomyces cerevisiae. Genetics. 159:441–452.

Wu J, Mizuno H, Hayashi-Tsugane M, Ito Y, Chiden Y, Fujisawa M, Katagiri S, Saji S, Yoshiki S, Karasawa W, et al. 2003. Physical maps and recombination frequency of six rice chromosomes. Plant J. 36:720–730. Doi:10.1046/j.1365-313X.2003.01903.x.

Yelina N, Diaz P, Lambing C, Henderson IR. 2015. Epigenetic control of meiotic recombination in plants. Sci China Life Sci. 58:223–231. Doi:10.1007/s11427-015-4811-x.

Zhu Y, Yin Y, Yang K, Li J, Sang Y, Huang L, Fan S. 2015. Construction of a high-density genetic map using specific length amplified fragment markers and identification of a quantitative trait locus for anthracnose resistance in walnut (*Juglans regia* L.). BMC Genomics. 16:614.

